# Regulation of virion production by the ORF8 signal peptide across SARS-CoV-2 variants

**DOI:** 10.1101/2024.03.05.583578

**Authors:** Mir M. Khalid, Irene P. Chen, Frank S. Soveg, Taha Y. Taha, Takako Tabata, Rahul K. Suryawanshi, Abdullah M. Syed, Alison Ciling, Maria McCavitt-Malvido, Ursula Schulze-Gahmen, Jennifer Hayashi, Ik-Jung Kim, Siew Wai Fong, Jyoti Batra, G. Renuka Kumar, Laurent Renia, Lisa FP Ng, Nevan J. Krogan, Jennifer A. Doudna, Eric Verdin, Melanie Ott

## Abstract

The open reading frame 8 (ORF8), an accessory protein of SARS-CoV-2, is prone to deletions and mutations across different viral variants, which was first described in several Singapore variants. The reason why viral evolution favors loss or inactivation of ORF8 is not fully understood, although the effects of ORF8 on inflammation, immune evasion, and disease severity have been described. Here we show –using clinical ORF8-deficient viral isolates, virus-like particles (VLPs) and viral replicons– that ORF8 expression dampens viral particle production. ORF8 physically interacts with the viral Spike protein and induces Golgi fragmentation, overall contributing to less virus particle production. Using systematic ORF8 deletions, we mapped the particle-reducing function to its N-terminal signal peptide. Interestingly, this part of ORF8 is severely truncated in the recent XBB.1.5 variant, and when restored, suppresses viral particle production in the context of the entire viral genome. Collectively, our data supports the model that evolutionary pressure exists to delete ORF8 sequence and expression across SARS-CoV-2 variants to fully enable viral particle production.

**Importance:** SARS-CoV-2 variants continue to emerge worldwide with advantages in replication and immune evasion. Many variants have acquired distinct mutations in independent lineages to abolish ORF8 expression. To understand the molecular mechanisms behind this evolutionary trend, we utilized reverse genetics, molecular virology, and confocal microscopy to show that ORF8 has antiviral functions by dampening viral particle production and inducing Golgi stress during infection. Our data demonstrate that SARS-CoV-2 is continuing its adaptation to optimize viral particle production and other unknown aspects of viral infection.

## Introduction

Since the beginning of the COVID-19 pandemic, numerous SARS-CoV-2 variants have and continue evolving with enhanced spread and antibody neutralization escape. The SARS-CoV-2 Spike protein has evolved the most with mutations altering its entry and immune evasion functions. Open reading frame 8 (ORF8) is a SARS-CoV-2 accessory protein that has also undergone continuous evolution since the beginning of the pandemic (**Fig S1**) and is often truncated in emerging viral variants without an apparent decrease in viral fitness (1–4). One such viral variant is an isolate from Singapore, where a 382-nucleotide deletion removed almost the entire ORF8 sequence as well as the 3’-end of ORF7b (5). Many other ORF8 mutations are common in circulating SARS-CoV-2 variants. Mutation S84L is now considered a lineage-defining mutation that is fixed in all circulating SARS-CoV-2, including all major variants of concerns (VOCs) and variants of interest (VOIs) (6) (**Fig S1**). The Alpha variant featured Q27Stop leading to the expression of a truncated protein (4,7). Similarly, a recent Omicron subvariant XBB.1.5 has an even shorter truncated ORF8 due to G8Stop mutation, which has been observed in all XBB.1.5-descendent variants including XBB.1.16, XBB.1.16, EG.1, EG.5 (4). Despite the many ways ORF8 expression is eliminated in SARS-CoV-2 variants, there was no apparent decrease in viral fitness as some of the variants have dominated globally. Hence, intact ORF8 might be deleterious to certain aspects of the viral life cycle.

Many studies have demonstrated that ORF8 is involved in immune modulation thereby likely being pro-viral. For example, ORF8 downregulates MHC-I expression and interferon responses (8,9). A characteristic feature of ORF8 is that it is secreted from infected cells and is readily detected in COVID-19 patient sera (1,10). As such, the protein has pro-inflammatory properties and binds for example to the IL17 receptor A (IL17RA), which might harm the host and thus suppress viral replication in the long-term (5,11–15). Overall, the role of ORF8 during viral replication is pleiotropic and not yet fully understood.

In the cell, ORF8 is an ER luminal protein, which binds and colocalizes with the Spike protein when co-expressed (16,17). ORF8 must transit through the Golgi for proper modification and secretion with the viral structural proteins that are also secreted. Structural proteins during this transit assemble into a viral particle and leave the cell as virions, while ORF8 has been shown to form dimers, adopting an immunoglobulin-like structure with an N-terminal 15-amino acid signal peptide that allows entry into the ER lumen (16,18,19). Once secreted, ORF8 elicits proinflammatory reactions through multiple host protein interactions, including binding to the IL-17RA (12,14,20). Here, we propose that its transit through the secretory pathway in the presence of the viral structural proteins, specifically Spike, allows ORF8 to regulate viral particle production directly. We test this hypothesis with virus-like particles (VLPs) that resemble authentic viral particles containing all four viral structural proteins packaging cargo genomes containing a SARS-CoV-2 packaging signal (21). Using this tool and reverse genetics, we investigated the impact of ORF8 on virion production.

## Results

### ORF8 expression reduces virion production but not genome replication

First, we compared the infectious virion production of the ancestral isolate WA1, which does not contain mutations in ORF8, and of the Singapore Isolate (abbreviated Sing), which lacks most of the ORF8 sequence (del27849_28230 (ORF8Δaa1-112)). We found that the Singapore isolate produces markedly more infectious virions than the WA1 isolate, supporting a model where ORF8 negatively regulates the viral life cycle (**Fig.1A**). To determine whether viral RNA replication is specifically involved, we turned to a replicon system based on the WA1 genome in which we replaced the Spike coding region with a secreted nano-luciferase (nLuc) reporter and eGFP (22). The lack of Spike prevents production of infectious virus upon infection. We generated a replicon lacking ORF8 expression by inserting a stop codon at the second amino acid (Rep-ORF8-Stop) (**Fig. 1B**). We transfected BHK-21 cells with either replicon plasmids along with a nucleocapsid protein (N) expression vector necessary to launch viral replication (22–25) (22–25) (**Fig.1B and Fig. S2B**). As part of viral replication, the Spike sgRNA will instead produce nLuc that is secreted into the supernatant. Therefore, luminescence from the cell culture media serves as a proxy for viral RNA replication as has been validated previously (22). We did not observe any difference in luminescence between the WA1 and the Rep-ORF8-Stop replicons (**Fig.1B**), indicating that ORF8 expression does not affect RNA replication.

**Figure 1:**
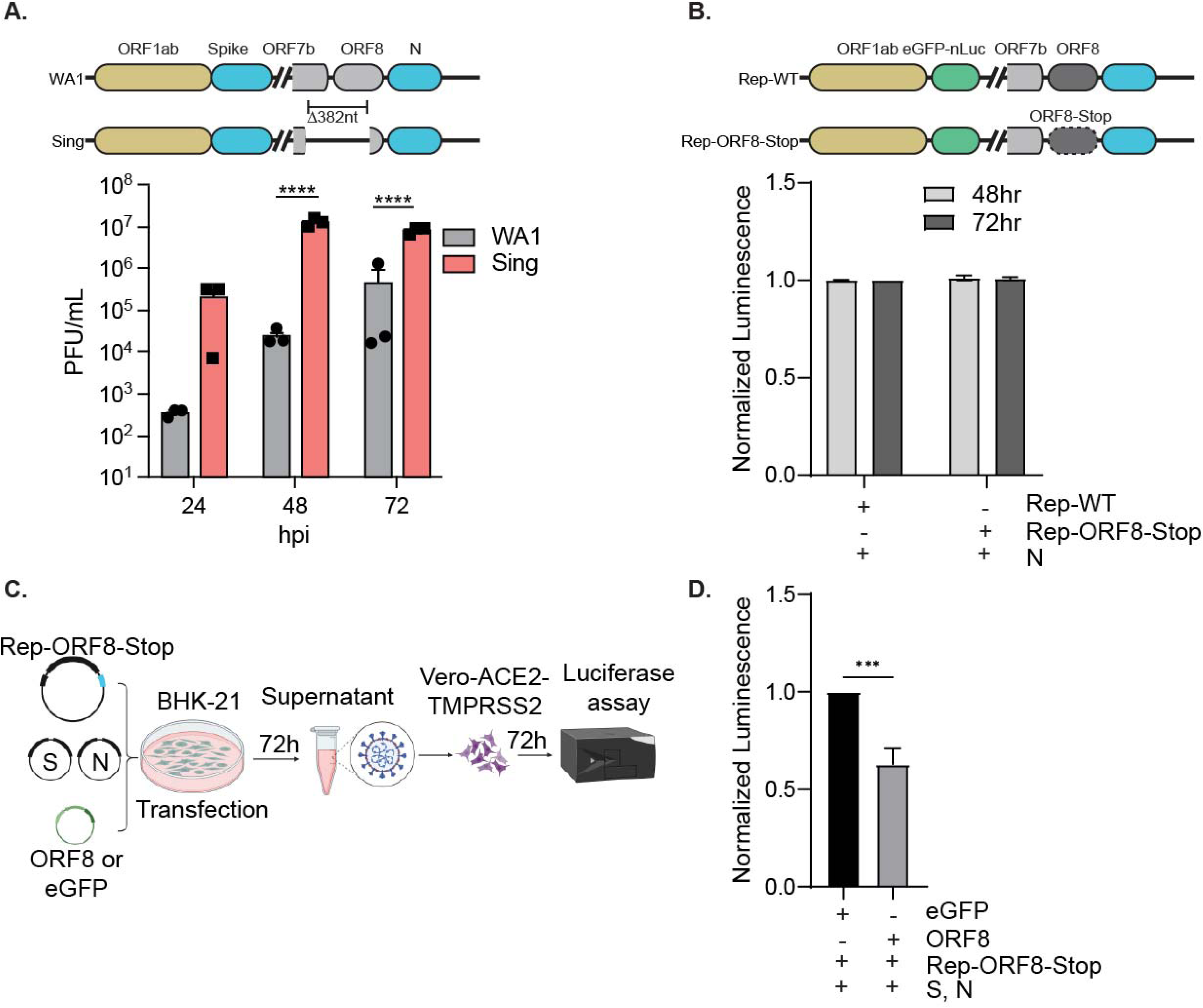
SARS-CoV-2 ORF8 reduces virus production but not genome replication. **A.** Top: Schematic diagram of WA1 (BEI NR-52281) and Singapore clinical isolate (EPI_ISL_414378). The Singapore isolate has a 382 nt deletion, comprising the 3’ end of ORF7b and removing almost all of the ORF8. Bottom: PFU assay of WA1 and Singapore isolate in HEK293T-ACE2-TMPRSS2 (HEK293T-A/T) cells. hpi: hours post-infection. **B.** Top: Schematic diagram of SARS-CoV-2 Spike-deleted replicon system. In Rep-WT, the whole SARS-CoV-2 genome except the Spike region was cloned into the pBAC vector using a pGLUE method previously developed by our lab (22). Spike is replaced with eGFP-nLuc, which is a reporter for genome replication in producer cells and infection in infected cells. The nLuc(Nano Luciferase) has a secretion signal peptide, which allows quantification of replication or infection from the supernatant. In Rep-ORF8-Stop, a stop codon at the second amino acid (K2Stop) prevents ORF8 synthesis. Bottom: genome replication for Rep-WT and Rep-ORF8-Stop plasmids. Luminescence of cell culture supernatant at different time points, normalized to Rep-WT condition at 48h time point. BHK-21 cells were transfected with different replicon plasmids in the presence of the Nucleocapsid (N) of SARS-CoV-2 Delta variant. **C.** Workflow for replicon production and infection experiment. BHK-21 cells are transfected with a replicon plasmid, plasmids expressing Spike and N from the SARS-CoV-2 Delta variant, and a plasmid expressing Strep-tagged eGFP (control) or Strep-tagged ORF8. Supernatants are collected at 72hr time points and used to infect Vero-ACE2-TMPRSS2 cells, later the supernatant of the infected cells was used for luminescence. Cell lysates are collected to confirm protein expression by western blot**. D.** Luminescence of culture media of replicon-infected Vero-ACE2-TMPRSS2 cells at 72 hpi. For each independent experiment, luminescence was normalized to the eGFP control condition. For each condition, three technical replicates were done for luminescence measurements with total 3 independent experiments were done. Data are represented as means plus SD. The P values were calculated by One-way ANOVA test (**B**), Two-way ANOVA test and two tailed paired t test (**D**). *, P ≤ 0.05; **, P ≤ 0.01; ***, P ≤ 0.001. n = 3 independent experiments.

Next, we repeated the assay but included a Spike expression vector in the transfection to produce single-round infectious viral particles. These particles can be used to perform single round infection in Vero ACE2 and TMPRSS2 cells. Luminescence signal after infection indicate the infectious titer of the generated single-round infectious particles. To investigate the effect of ORF8 expression on particle production, we used the Rep-ORF8-Stop construct and co-expressed Strep-tagged ORF8 -or eGFP control plasmid along with the Spike and N expression vectors (**Fig. 1C**). We found that ORF8 expression in producer cells significantly reduced the production of single-round infectious particles (**Fig. 1D**). We conclude that ORF8 has a negative impact on particle production or entry but does not affect viral RNA replication.

### ORF8 colocalizes with Spike in Cis-Golgi and causes Golgi stress response in infected cells

During infection, SARS-CoV-2 structural proteins S, N, M, and E as well as genomic RNA assemble at the ER-Golgi intermediate compartment (ERGIC) to produce infectious virions. We previously showed that ORF8 is an ER-luminal protein and interacts with Spike when overexpressed (16). To determine whether virally produced ORF8 colocalizes with Spike during infection, we infected A549-ACE2 cells with the WA1 and Singapore clinical isolates, and examined ORF8 and Spike localization relative to GM130, a cis-Golgi marker. ORF8 and Spike colocalized with GM130 only in WA1-infected cells (**Fig. 2 and Fig. S3**). In addition, we observed Golgi fragmentation and dispersion in WA1-infected cells, whereas in mock- and Singapore-infected cells, the GM130 signal showed a normal dotted pattern near the nucleus, despite high viral particle production in the latter case. This indicates that ORF8 compromises the integrity of the cellular Golgi apparatus similarly to what was previously observed upon SARS-CoV-2, flavivirus infection, and flavivirus NS1 protein expression (26–28). These data indicate that ORF8 colocalizes with Spike in intracellular ER-Golgi compartments during infection, which appear disrupted and fragmented over time only when ORF8 is present.

**Figure 2:**
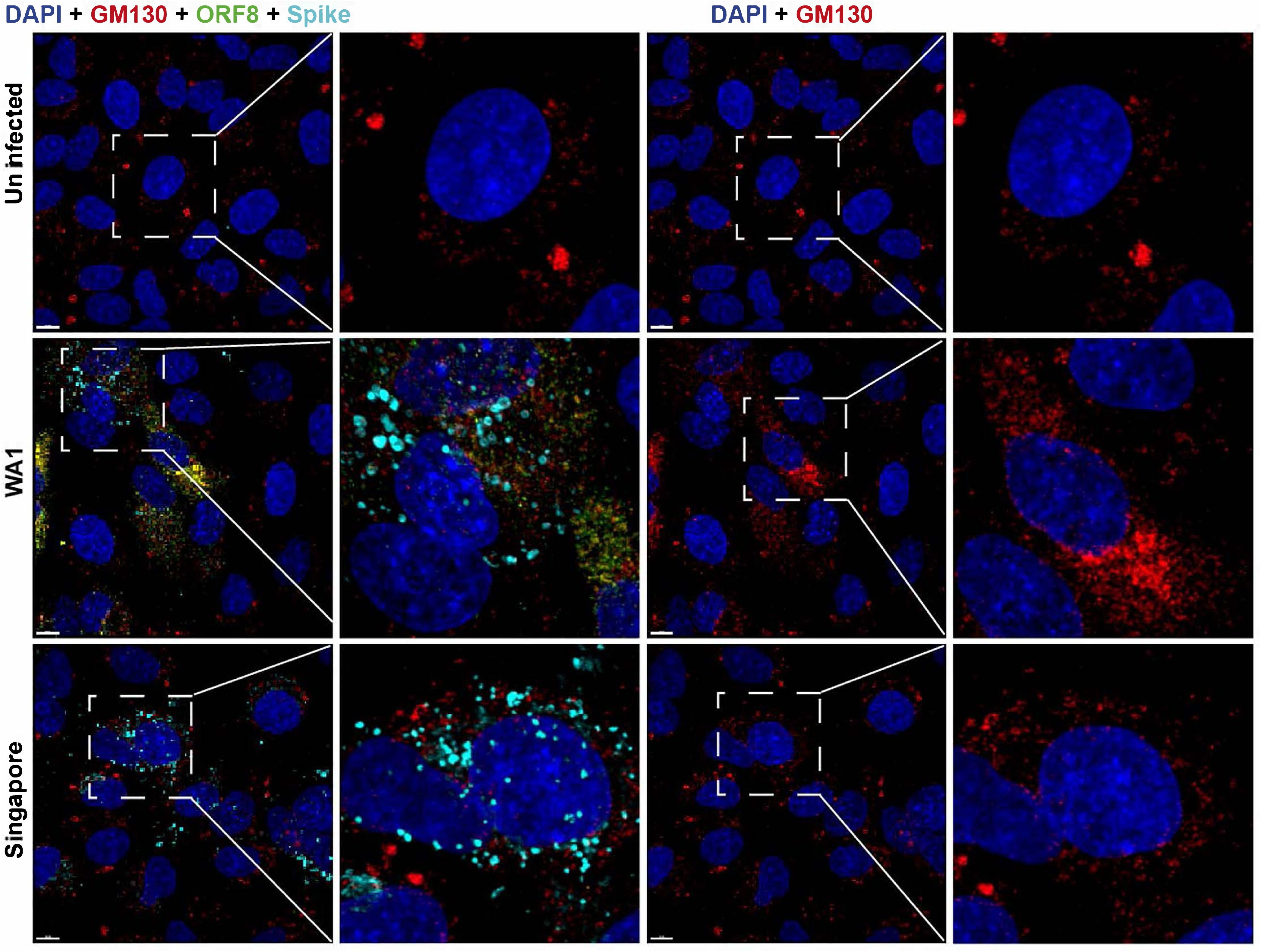
SARS-CoV-2 ORF8 co-localizes with Spike in cis-Golgi (GM130) and induces stress in Golgi (appears disrupted and fragmented). A549-ACE2 cells were infected with WA1 (BEI NR-52281) and Singapore (EPI_ISL_414378) clinical isolates. Immunostaining was done with antibodies against ORF8 (Green), Spike (Turquoise) and the cis-Golgi marker GM130 (Red). DAPI (Blue) was used to stain nuclei. White dotted square area was enhanced three times. White bar = 8 μm. n = 3 independent experiments.

### ORF8 reduces VLP production but not VLP entry

To further define the effect of ORF8 on virion production, we generated VLPs in the presence or absence of ORF8. Similar to replicon particles, VLPs are an authentic tool for investigating SARS-CoV-2 viral particle production and entry (21,29) as they also contain the four viral structural proteins S, N, M, and E but instead of a modified or intact viral genome, they package a reporter RNA with containing the packaging sequence, named T20 (21). To this end, we transfected HEK-293T cells (producer cells) with plasmids expressing SARS-CoV-2 S, N, M, and E proteins, a reporter plasmid containing the T20 packaging signal and firefly luciferase and a plasmid expressing either eGFP or ORF8 (**Fig. 3A**). Once expressed in HEK-293T cells, the structural proteins assemble VLPs that contain the T20 reporter RNA and are released into the culture medium. These VLPs are then used to infect HEK293T-ACE2-TMPRSS2 (receiver cells). The luminescence from infected cells is a measurement of the VLP titer (21). In a dose-dependent manner, we confirmed that VLPs produced in the presence of ORF8 show less luminescence in infected cells than VLPs produced in the absence of ORF8 (**Fig. 3B**).

**Figure 3:**
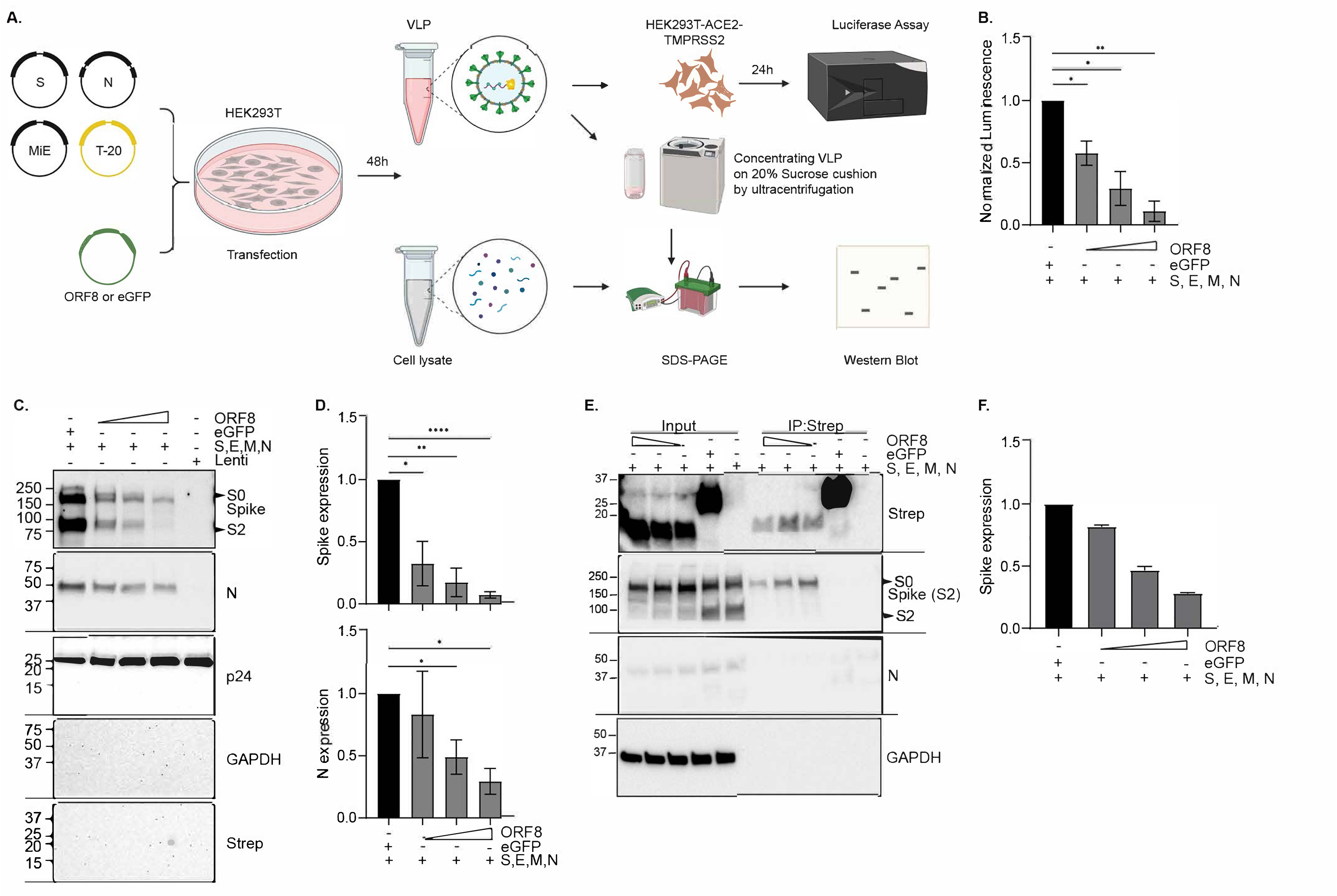
SARS-CoV-2 ORF8 expression reduces the production of virus-like particles. **A.** Schematic representation of virus-like particle (VLP) production, confirmation, and quantification. Plasmids encoding SARS-CoV-2 (B.1 lineage) structural proteins (S(Spike), N (nucleocapsid protein), MiE (membrane and envelope, expressed from the same plasmid), and the SARS-CoV-2 packaging signal with firefly luciferase reporter (T-20) were transfected in HEK293T cells, along with either a plasmid encoding a Strep-tagged eGFP (control), or three different doses of a plasmid encoding a Strep-tagged ORF8. VLP-containing culture media was collected 48 hr after transfection. This VLP was either used to infect HEK293T-ACE2-TMPRSS2 cells (HEK293T-A/T) or purified on 20% sucrose cushion for further analysis. For infection in HEK293T-A/T (VLP receiving cells), luminescence was measured from the cell 24 hr later. Cell lysates were collected from transfected HEK293T cells (VLP producer cells). Both cell lysates, and purified VLP were ran on SDS-PAGE gel to assess protein expression by western blot. **B.** Luminescence readout of HEK293T-A/T cells infected with VLPs produced in the presence of increasing amounts ORF8, normalized to the eGFP control. For each condition, three technical replicates were done for luminescence measurements with total 3 independent experiments were done. **C.** Western blot analysis of proteins in the concentrated VLP fractions. P24 was added to the VLPs prior to sucrose purification as a control for concentration. **D.** Spike and N amounts in concentrated VLP particles as a function of ORF8 plasmid amounts, normalized to the eGFP control condition. Protein amounts were derived from densitometry analysis of western blots. **E.** Western blot analysis of HEK293T cells lysates (Input) and co-immunoprecipitation by Strep pull down (IP:Strep). For immunoprecipitation, the same amount of cell lysate across all conditions was pulled down with streptactin sepharose resin. **F.** Quantification of Spike in the HEK293T cells lysates as a function of ORF8 plasmid dose. Spike abundance was measured by densitometry analysis of western blots and normalized to the eGFP control condition. Data are represented as means plus SD. The P values were calculated by One-way ANOVA test. *, P ≤ 0.05; **, P ≤ 0.01; ***, P ≤ 0.001; ns, not significant. n = 3 independent experiments

This can be interpreted as less cellular entry of the VLPs into the receiver cells or a reduction in producing infectious particles. We, therefore, purified VLPs produced either in the presence or absence of ORF8 on a 20% sucrose cushion and measured the proteins contained in this fraction by western blot (**Fig. 3A**). By comparing the amount of Spike and N proteins on the particles, we found that ORF8 expression leads to a significant reduction of Spike protein loading and to a lesser extent of N protein content in the secreted particles (**Fig. 3C-D).** This indicates that VLP production is reduced when ORF8 is present in producer cells.

To examine further the possibility that virion entry is affected by ORF8, we generated VLPs without ORF8 and incubated them with recombinant ORF8 before infecting the receiver cells. We saw no difference in the receiver cells’ luminescence or viability (**Fig.S4A**), indicating that recombinant ORF8 does not interfere with VLP entry into susceptible cells. We conclude that ORF8 interferes with VLP production in producer cells but not entry into receiver cells.

We next asked whether ORF8 interacted with the structural proteins within VLP producing cells. Co-immunoprecipitation experiments confirmed that ORF8 interacted with the Spike but not with N in the VLP-producing cells (**Fig. 3E**). Due to antibody issues, we could not determine whether ORF8 interacts with M and E during VLP production. Notably, our analysis of the producer cells’ lysates also showed that ORF8 expression decreased Spike expression in a dose-dependent manner in co-expressing cells (**Fig. 3F**). This was also observed when we introduced 4-fold more of one of the structural proteins (S, N, M, and E plasmids) together with steady amounts of ORF8 into cells to exclude promoter competition or technical issues as an explanation (**Fig.S4 B-D**). These findings confirm previous data that ORF8 reduces Spike protein expression and show newly that this happens in VLP-producing cells independently of plasmid amount or promoter competition.

Given that SARS-CoV-2 VLP production depends on specific Spike concentrations, we can conclude that ORF8 reduces total VLP production (21). Notably, we could not detect ORF8 in purified particles. This does not exclude the possibility that ORF8 can bind Spike on virus particles. Together, these findings suggest that ORF8 specifically reduces Spike amounts in producer cells, thus hindering VLP production and causing the release of fewer VLPs.

### The ORF8 signal peptide (aa 1–15) is sufficient to reduce VLP production

We tested whether the recent variants Delta or Omicron(BA.1) can escape ORF8-mediated reduction in VLP production. Interestingly, we saw ORF8 efficiently reduced VLP production in these variants **(Fig.S5).** In addition, we determined by site-directed mutagenesis that none of the seven cysteine (C) residues that cause intermolecular (C20) and intramolecular (C25 with C90, C37 with C102 and C61 with C83) disulfide bonds within ORF8 are required for the Spike-suppressive activity of ORF8 (**Fig.S6**).

To find the minimal region of ORF8 responsible for reducing VLP production, we divided ORF8 into three distinct regions (aa 1–40, aa 41–80, and aa81–121), which we cloned into an eGFP plasmid to produce eGFP-ORF8 fusion proteins. We also cloned aa 1–7 and aa 1–26 of ORF8 into the eGFP plasmid, mimicking ORF8 from XBB.1.5 (G8Stop) and Alpha (Q27Stop) variants, respectively. We generated VLP in the presence of these eGFP-ORF8 constructs. All constructs reduced VLPs production, relative to the eGFP control plasmid, except for the aa 41-80 construct, which behaved essentially like the eGFP control (**Fig. 4A**). Interestingly, having the first 26 aa was enough to reduce VLP production nearly to the same extent as the full-length ORF8, while having the first 7 aa reduced it by 50% (**Fig. 4A**). The fragments’ effect on VLP production mostly correlated with their ability to bind the Spike protein: we saw little to no interaction between Spike and aa 1–7 or aa 41–80, but a strong interaction with aa 1–26 and aa 1–40 (**Fig.S7A**). This observation indicates that the first 26 aa of ORF8 are sufficient to bind Spike and can cause a reduction in VLP production. Nevertheless, Spike interaction with ORF8 is not necessary for reducing VLP production since aa 81–121 strongly reduced VLP production even though it did not bind Spike. This last finding underscores that at least two regions of ORF8 can inhibit VLP production: an N-terminal portion that binds Spike, and a C-terminal portion that does not. Removal of 40 aa from the N-terminal, middle, and C-terminal portions of ORF8 confirmed that only the N-terminal portion is required to bind Spike (**Fig. 4B, S7B**). Since Alpha ORF8 has a stop codon at position 27 in the N terminus, it still binds Spike, but the lack of the C-terminal region reduces its additional suppressive effect on VLP production.

**Figure 4:**
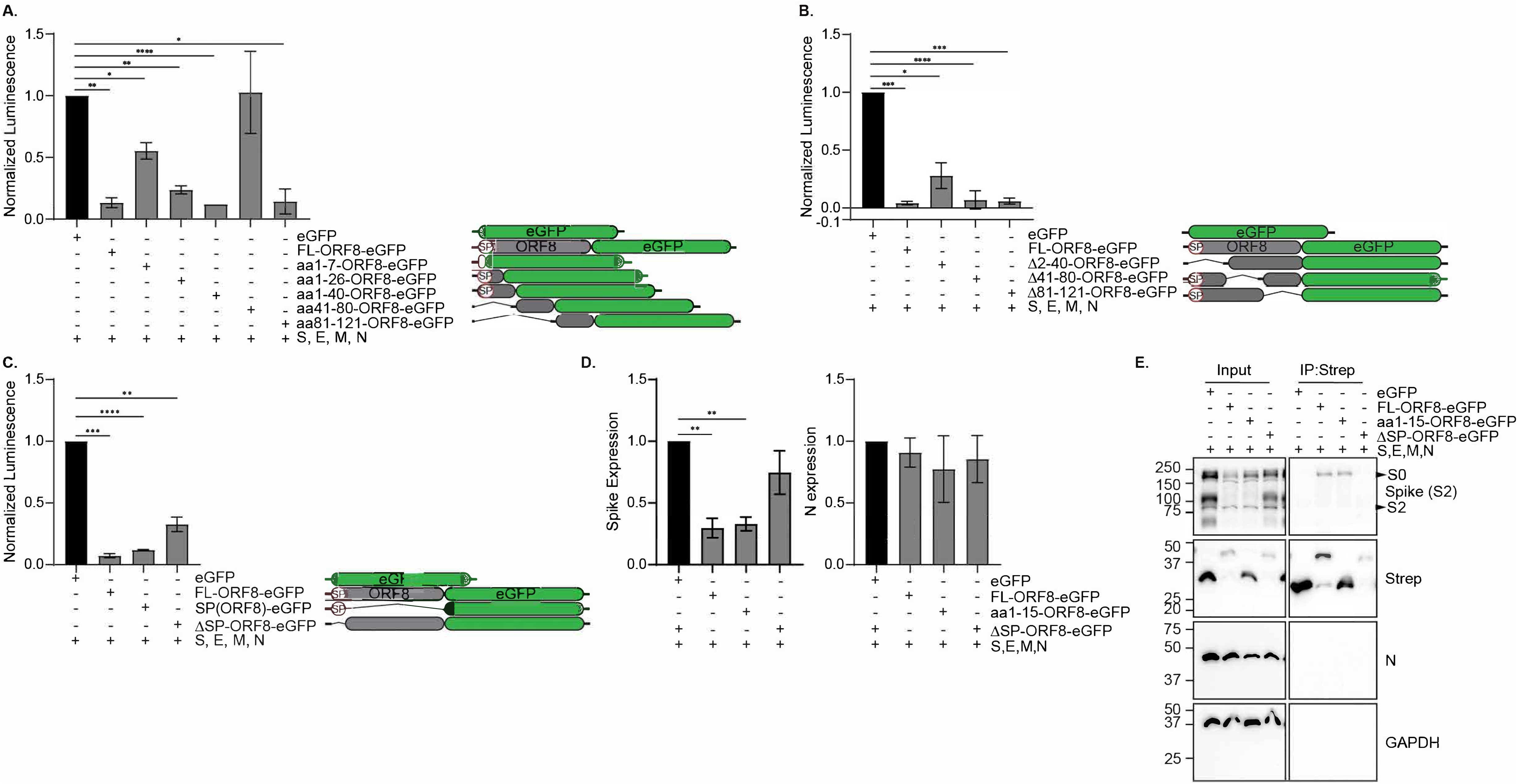
Expression of the N-terminal 15 aa signal peptide of ORF8 reduces SARS-CoV-2 VLP production. **A -C.** Normalized luminescence of VLP-infected cells. VLPs were produced in the presence of different fragments of ORF8. The ORF8 fragments were expressed as fusions to eGFP. Luminescence is normalized to the eGFP control condition in each experiment. For each condition, three technical replicates were done for luminescence measurements with total 3 independent experiments were done. FL = Full Length, SP = Signal peptide. **D.** Expression of Spike and N protein in VLP-producing cells. Densitometry of western blots was used for protein quantification. **E**. Representative western blot of VLP-producing cells. Immunoprecipitation was done using Strep pulldown. Data are represented as means plus SD. The P values were calculated by One-way ANOVA test. *, P ≤ 0.05; **, P ≤ 0.01; ***, P ≤ 0.001; ns, not significant. n = 3 independent experiments.

ORF8 has a 15 aa signal peptide at its N-terminus. Based on our data with the truncated ORF8 of the XBB.1.5 and Alpha variants, we speculated that the signal peptide might play a role in reducing VLP production. We therefore generated a construct that only expresses the signal peptide of ORF8 and a construct that expresses ORF8 without its signal peptide and tested their impact on VLP production. The signal peptide alone significantly reduced VLP production (**Fig. 4C)**. The ORF8 devoid of signal peptide allowed more VLP production but did not fully rescue VLP production, presumably because it retained the inhibitory C-terminal region of ORF8. We also found that the ORF8 signal peptide reduced Spike levels and interacted with Spike, while the ORF8 lacking the signal peptide did not (**Fig. 4D-E**). We performed a triple alanine scan from aa 16 to 42 in ORF8 and found no changes in VLP production or interaction with Spike (**Fig. S8A-C**). We conclude that the 15-aa ORF8 signal peptide is sufficient to bind Spike and reduce VLP production.

### Restoration of ORF8 in XBB.1.5 reduces virus and VLP production

The recently emerged XBB.1.5, XBB.1.16, EG.1, and EG.5 lineages share the G8Stop mutation in ORF8. To determine the relevance of the stop codon, we compared XBB.1.5, and XBB.1.16 VLP production in the presence of full-length and truncated (aa 1-7) ORF8, compared to eGFP. We saw a significant reduction in VLP production with full-length ORF8 but not with the truncated version, as expected (**Fig.5A**).

**Figure 5:**
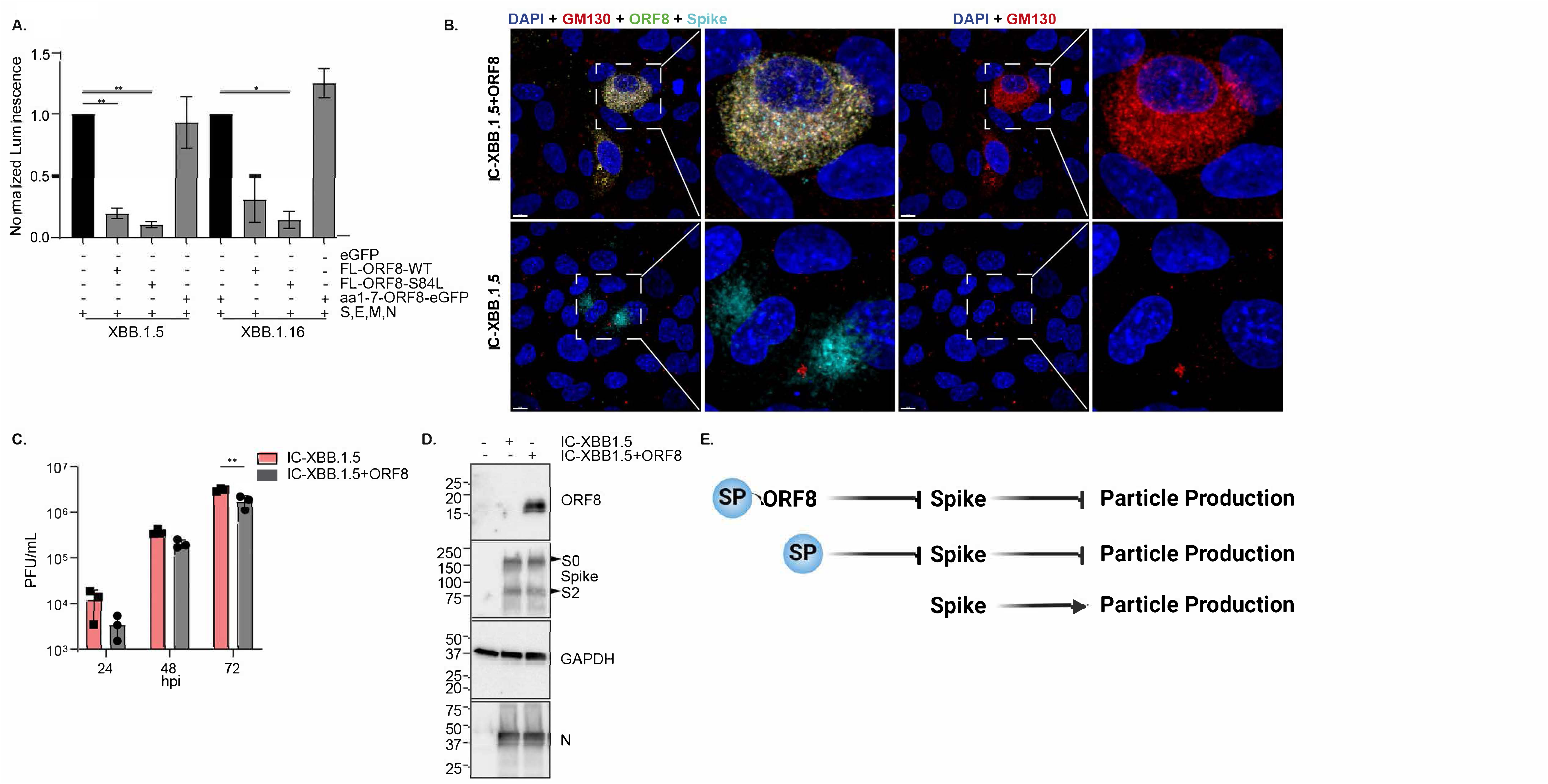
Truncated ORF8 from XBB.1 variants increases VLP production. **A.** Normalized luminescence of cells infected with VLPs from various XBB variants. VLPs were produced in the presence of different ORF8s. For each condition, three technical replicates were done for luminescence measurements with total 3 independent experiments were done. **B.** A549-ACE2 cells were infected with recombinant XBB. IC-XBB.1.5 (contains naturally occurring ORF8-G8Stop mutation) and IC-XBB.1.5+ORF8 (ORF8 full-length expression was restored by restoring Glycine at amino acid 8 position in ORF8). Immunostaining was done with antibodies against ORF8 (Green), Spike (Turquoise) and the cis-Golgi marker GM130 (Red). DAPI (Blue) was used to stain nuclei. White dotted square area was enhanced three times. White bar = 8 μm. **C.** Plaque forming assay in Calu3 cells infected with the recombinant XBB.1.5 virus. **D.** Representative western blot of cells infected with recombinant XBB.1.5 virus. **E.** A model summarizing the role of ORF8 hampering SARS-CoV-2 viral particle production. In the presence of ORF8, ORF8 interacts with Spike, reducing its expression. We also observed Golgi fragmentation and dispersion in isolates that express ORF8. This led to disrupted viral particle production, reducing viral particle production. The signal peptide sequence alone is enough to interact with spikes and reduce spike expression and viral particle production. Without ORF8, all the structural proteins are properly expressed, and particle production is not hampered. We also observed intact Golgi in infected cells by ORF8 deleted or truncated isolates. Thus, it leads to increased or normal levels of viral particle generation. Data are represented as means plus SD. The P values were calculated by One-way **(A)** ANOVA test, and Two-way (**C**) ANOVA test. *, P ≤ 0.05; **, P ≤ 0.01; ***, P ≤ 0.001. n = 3 independent experiments.

To further prove the importance of the stop codon for viral fitness, we restored ORF8 expression in XBB.1.5 by converting the stop codon to Glycine (we refer to this virus as IC-XBB.1.5+ORF8). In A549-ACE2 cells infected by this virus, we saw that ORF8 colocalized with Spike and GM130, like we previously observed (**Fig.5B**). We also observed fragmented Golgi morphology upon infection with IC-XBB.1.5+ORF8 but not by XBB.1.5 (have truncated ORF8). When we infected Calu3 cells with both viruses, we saw a significant reduction in particle production with IC-XBB.1.5+ORF8 virus at 72 hpi (**Fig.5C**). ORF8 expression in IC-XBB.1.5+ORF8 virus-infected cells was confirmed (**Fig.5D**). These data indicate that the introduction of a stop codon at position eight as seen in the XBB.1.5 variant is beneficial for infectious virus production. This supports our model that ORF8 expression dampens viral particle production that deletions and mutations affecting the signal peptide of ORF8 are evolutionary events that counterbalance this suppression (**Fig.5E**).

## Discussion

Most studies so far have focused on the immunomodulatory and host-response function of ORF8, less is known about the role ORF8 may play in the SARS-CoV-2 replication cycle. We have found that the signal peptide and the C terminal domain of ORF8 can reduce SARS-CoV-2 virus production by hampering viral particle production. Notably, we did not find any effects of ORF8 on entry and RNA replication steps. Our work confirms findings from previous studies, including our own, that described an interaction between Spike and ORF8, a reduction in Spike pseudovirus particle productions, but also, importantly, extends our knowledge by showing ORF8 targeting viral particle productions and causing Golgi fragmentations using live virus, replicon, and VLP. While we contribute significant knowledge in VLP, replicon and virus-infected cells, our study also opens several new areas of study.

Our model is that in SARS-CoV-2, which has intact ORF8 or produces signal peptides of ORF8, viral particle production is dampened, possibly as a way to reduce immunosurveillance of infected cells (14,16,30) or as an unwanted effect of ORF8’s other functions benefitting viral replication (8,9,18) (**Fig 5E)**. In ORF8-deficient SARS-CoV-2, viral particle production is not disrupted, leading to increased viral particle formation (**Fig 5E)**. The effect on viral particle production happens through ORF8 binding Spike by its signal peptide. We also found that ORF8 expressions in infected cells can cause Golgi fragmentation.

In case of flavivirus infection, Dengue virus (DENV) NS1 protein can cause Golgi stress as presented with Golgi dispersion and fragmentation (27). DENV NS1 also has a 24 aa signal peptide sequence, forms dimer, and gets secreted as a soluble protein (31,32). ORF8 signal peptide may be behaving in a similar way as NS1. Like NS1, ORF8 can be secreted by conventional and unconventional secretory pathways (14,33). Golgi morphology can change when Golgi stress response is activated to cope with cells’ capacity to mature, traffic, and secrete proteins using the Golgi network (34). Secreted proteins cannot get secreted or delivered to the plasma membrane if Golgi traffic is disrupted. Previously we saw reduced Spike co-localization in the trans-Golgi network along with less Spike antigen on cells, when ORF8 and Spike are co-expressed (16). It is plausible that ORF8 by itself or with other proteins, causes Golgi stress, reducing Spike availability to be incorporated in viral particles, competing with virus for its own secretion-hence reducing virus particles to form and exit out of infected cells.

At the same time, secreted ORF8 causes cytokine storms and activation of inflammatory pathways (14,15). Patients vaccinated with inactivated SARS-CoV-2 have anti-ORF8 antibodies in their serum-suggesting that the secreted ORF8 (probably in the inactivated vaccine) can activate immune responses (30). Thus, the secretory nature of ORF8 can bring more attention to virus-producing cells.

Many SARS-CoV-2 variants encode a truncated version of ORF8 or limit its expression. The Singapore variant has almost a deleted ORF8, and it has been linked to reduced severity (5). Similar isolates have been reported in other parts of the world, such as Australia, Taiwan, and Bangladesh (35). Although these isolates did not emerge as major variants, their presence shows the continuous evolution of SARS-CoV-2. In recent lineages, ORF8 deletion or truncations are present in many sequences (3,4) (**Fig. S1**). For sequences with full-length ORF8, such as BA.5 and XBC, it has been observed that transcriptional regulatory sequences (TRS) contain a mutation that is predicted to reduce ORF8 expression. Hisner et al. summarized and analyzed recent variants with these ORF8 TRS mutations. They predicted that ORF8 ablation would dominate in future lineages, while current lineages show reduced or removed ORF8 expression (4). The XBB.1.5, XBB.16, and EG.5 lineages express a truncated ORF8, only the N-term 7 amino acids. In addition, the recent emergence of the BA.3.2 lineage with an 870-nt deletion removing ORF7a and ORF8, further supports the relevance of investigating ORF8 function and deletion events to better understand SARS-CoV-2 adaptation and COVID-19.(36,37)

It is interesting to see that ORF8 is intact in its natural host. Comparing ORF8 of Bat coronavirus (RaTG13) with SARS-CoV-2 ORF8 shows different interaction with monocytes (38), suggesting a potential role for ORF8 in host adaptation. With these truncations and deletions, it is plausible that SARS-CoV-2 evolution shapes minimal ORF8 length for optimal viral replication in humans.

### Limitations of the study

Due to antibody issues, we could not determine whether ORF8 might have some roles in the other two structural proteins (M and E) during VLP production, but we successfully excluded interactions with N. While we show a role of ORF8 in viral particle production, we are currently distinguishing between possible roles in viral assembly and egress. We are keen to study the function of ORF8 in Golgi fragmentation and the role of this function on viral assembly in future studies. Further investigation on this can lead to an understanding of the biology of SARS-CoV-2 and the role of Golgi stress in viral replication.

## Methods

### Ethics

We performed all research following all relevant ethical regulations. All work conducted with replication-competent SARS-CoV-2 viruses was done in an approved biosafety level 3 (BSL3) laboratory and experiments approved by the Institutional Biosafety Committee of the University of California, San Francisco, and Gladstone Institutes. The VLP and replicon work was done in an approved biosafety level 2 (BSL2) laboratory.

### Virus

SARS-CoV-2 USA-WA1/2020 (BEI NR-52281) and Singapore (EPI_ISL_414378) clinical isolates were used for all live virus infection studies. All live virus experiments were performed in a Biosafety Level 3 laboratory. The virus stocks were propagated in Vero-E6 cells, and their sequence was verified by next-generation sequencing. Viral stock titer was calculated using plaque-forming assays.

### Infectious clone and replicon generation

Rep-WT, Rep-ORF8-stop, icXBB.1.5, and icXBB.1.5+ORF8 were generated following pGLUE method in the pBAC SARS-CoV-2 construct as previously described (22). Briefly, the SARS-CoV-2 genome was divided into 10 fragments, which were cloned and later ligated via BsaI digestion and Golden Gate assembly protocol as described by Taha et al (22,39). The ligated plasmid was confirmed via nanopore sequencing (Primordium Labs). For XBB.1.5 infectious clone generation, BHK cells were transfected with 3 μg of pBAC-SARS-COV-2 construct and1 μg of SARS-CoV-2 N (Delta) plasmid with the X-treme Gene 9 (X9) DNA transfection reagent (Roche: 6365809001) was added 1:3 as DNA:X9. The supernatant was collected after 3 days post-transfection and used to infect Vero ACE2 TMPRSS2 cells. The sup from infection was passaged further to achieve high titer. All viruses generated in this study were sequenced and verified by NGS. All infectious clone experiments were done in BSL3. For replicon production, BHK cells were transfected with pBAC-SARS-COV-2 construct (1), SARS-CoV-2 Spike (0.5), SARS-CoV-2 N (0.5), and ORF8(4) plasmid with indicated molar ratio with Xtreme gene 9. In experiments where eGFP replaced ORF8, eGFP amount was used half of ORF8 amount (ORF8 expression was lower than eGFP), and supplemented with vector plasmid. The medium was replaced 12-16 hours later and added fresh culture media. Medium-containing replicon was collected 48 and 72 hours later post-transfection. For infection, 100μl of replicon was used to infect Vero-ACE2-TMPRSS2 cells. The cells were plated in 3×10^4 cells/well in 96 well plate the day before infection. The day after infection, medium was replaced and 72 hours post-infection 50μl of sup was used for nano-luciferase assay (Promega, N1120) according to manufacturer protocol. Luminescence was measured using a TECAN plate reader (Spark) with 30s shaking, auto attenuation, and 1000 ms integration. At least 3 independent experiments and infections were done in three technical replicates.

### Cell lines

HEK293T, BHK-21, Calu3 and Vero-E6 were obtained from ATCC were cultured in DMEM (Corning) supplemented with 10% fetal bovine serum (FBS) (GeminiBio), 1% glutamine (Corning), and 1% penicillin-streptomycin (Corning) at 37°C, 5% CO2. Calu3 cells were cultured in Advanced (Gibco) supplemented with 2.5% FBS, 1% GlutaMax, and 1% penicillin-streptomycin at 37°C, 5% CO2. A549 cells stably expressing ACE2 (A549-ACE2) were generated by transducing with ACE2-encoding (generated using Addgene plasmid no. 154981, a gift from Sonja Best, Rocky Mountain Labs, Hamilton, MT, USA) lentiviruses and selection with blasticidin (10 µg/mL) for 10 days. ACE2 expression was verified by Western blot. A549-ACE2 cells were cultured in DMEM supplemented with 10% FBS, blasticidin (20 μg/ml) (Sigma), and maintained at 37°C with 5% CO2. The HEK-293T ACE2-TMPRSS2 cell line was generated using sequential transduction as described in (29). Vero stably coexpressing human ACE2 and TMPRSS2 cells (gifted from A. Creanga and B. Graham, NIH, Bethesda, MD) were cultured at 37 °C and 5% CO2 in DMEM (Gibco) supplemented with 10% fetal calf serum, 100 ug/mL penicillin and streptomycin (Gibco), and 10 μg/mL of puromycin (Gibco).

### Plaque forming assay

Cell supernatants were analyzed for viral production using plaque-forming assays. Vero ACE2 TMPRSS2 cells were used for the assay and plated 24 hr before infection. Then the supernatants from producer cell cultures were serially diluted in DMEM (Corning) and added on to the cells. After a 1 hr absorption period, 2.5% Avicel (Dupont, RC-591) was overlaid. The overlay was removed 72 hours later, fixed in 10% formalin for 1 hour, and stained with crystal violet for 10 minutes to visualize plaque formation. All plaque assays were done in two technical duplicates from 3 independent experiments.

### Plasmids

The plasmids expressing SARS-CoV-2 ORF8 (ORF8-WT) were generous gifts from Dr. Nevan Krogan (UCSF, The Gladstone Institutes). An ORF8-WT plasmid was used as a source for generating all the mutated plasmids. Most of the ORF8 mutants were generated by designing primers with the desired mutations, generating PCR fragments with these primers and ligating them. The rest of them were generated with custom gene fragments (gblock(IDT)). All these mutation-containing fragments, PCR products or gblock, were ligated using NEBuilder HiFi DNA Assembly Master Mix (E2621L). The ligated product was transformed to Mach1-competent cells. The plasmids expressing SARS-CoV-2 Spike(S), Nucleocapsid (N), Membrane (M), and Envelop (E) for VLP production were generous gifts from Prof. Jennifer A. Doudna (The Gladstone Institutes). All plasmids used in this study are listed in Table 1. All plasmids and corresponding sequence information are available upon request.

**Table 1.**

| **Plasmid name** | **content** | **Addgene No** |
| --- | --- | --- |
| pLVX-EF1a-IRES-Puro | Vector | N/A |
| eGFP-2xStrep | eGFP | 141395 |
| CoV2-Spike-D614G | B.1 Spike | 177960 |
| CoV2-Spike-Delta | Delta Spike | N/A |
| CoV2-Spike-Omicron | Omicron (BA.1) Spike | N/A |
| CoV2-Spike-XBB.1.5 | XBB.1.5 Spike | N/A |
| CoV2-Spike-XBB.1.16 | XBB.1.16 Spike | N/A |
| CoV2-N-B.1.1.7 | B.1 N | 177958 |
| CoV2-N-Delta | Delta N | N/A |
| CoV2-N-Omicron | Omicron (BA.1) N | N/A |
| CoV2-N-XBB.1.5 | XBB.1.5 N | N/A |
| CoV2-N-XBB.1.16 | XBB.1.5 N (Same mutations) | N/A |
| CoV2-M-IRES-E | B.1 (M, and E) | 177938 |
| CoV2-M-IRES-E-Delta | Delta (M, and E) | N/A |
| CoV2-M-IRES-E-Omicron | Omicron (BA.1) (M, and E) | N/A |
| CoV2-M-IRES-E-XBB.1.5 | XBB.1.5 (M, and E) | N/A |
| CoV2-M-IRES-E-XBB.1.16 | XBB.1.5 (M, and E) (Same mutation)s | N/A |
| Luc-T20 | Luciferase | 177941 |
| ORF8-FL-2Xstrep-eGFP | Full length ORF8 fused eGFP | N/A |
| ORF8-aa1-40-2Xstrep-eGFP | ORF8 aa2-40 fused eGFP | N/A |
| ORF8-aa41-80-2XStrep-eGFP | ORF8 aa41-80 fused eGFP | N/A |
| ORF8-aa41-121-2XStrep-eGFP | ORF8 aa41-121 fused eGFP | N/A |
| ORF8-Δ2-40-2Xstrep-eGFP | ORF8 except aa2-40 fused eGFP | N/A |
| ORF8-Δ41-80-2XStrep-eGFP | ORF8 except aa41-80 fused eGFP | N/A |
| ORF8-Δ41-121-2XStrep-eGFP | ORF8 except aa41-121 fused eGFP | N/A |
| ORF8-aa1-7-2Xstrep-eGFP | XBB.1.5 ORF8-fused eGFP | N/A |
| ORF8-aa1-26-2Xstrep-eGFP | Alpha ORF8-fused eGFP | N/A |
| ORF8-SP-2Xstrep-eGFP | ORF8 aa1-15 fused eGFP | N/A |
| ORF8-ΔSP-2Xstrep-eGFP | ORF8 except aa1-15 fused eGFP | N/A |
| ORF8-WT or ORF8-FL | Full length ORF8 | 141390 |
| ORF8-C20A-2XStrep | FL-ORF8 with aa20 changed to Alanine | N/A |
| ORF8-C25A-2XStrep | FL-ORF8 with aa25 changed to Alanine | N/A |
| ORF8-C37A-2XStrep | FL-ORF8 with aa37 changed to Alanine | N/A |
| ORF8-C61A-2XStrep | FL-ORF8 with aa61 changed to Alanine | N/A |
| ORF8-C83A-2XStrep | FL-ORF8 with aa83 changed to Alanine | N/A |
| ORF8-C90A-2XStrep | FL-ORF8 with aa90 changed to Alanine | N/A |
| ORF8-C102A-2XStrep | FL-ORF8 with aa102 changed to Alanine | N/A |
| ORF8-NoCys-2XStrep | FL-ORF8 with all cysteine changed to Alanine | N/A |
| ORF8-S84L-2XStrep | FL-ORF8 with aa84 changed to leucine | N/A |
| ORF8-AAA-16-18 | FL-ORF8 with triple Alanine substitution at amino acids 16-18 | N/A |
| ORF8-AAA-19-21 | FL-ORF8 with triple Alanine substitution at amino acids-AAA-19-21 | N/A |
| ORF8-AAA-22-24 | FL-ORF8 with triple Alanine substitution at amino acids-AAA-22-24 | N/A |
| ORF8-AAA-25-27 | FL-ORF8 with triple Alanine substitution at amino acids-AAA-25-27 | N/A |
| ORF8-AAA-28-30 | FL-ORF8 with triple Alanine substitution at amino acids-AAA-28-30 | N/A |
| ORF8-AAA-31-33 | FL-ORF8 with triple Alanine substitution at amino acids-AAA-31-33 | N/A |
| ORF8-AAA-34-36 | FL-ORF8 with triple Alanine substitution at amino acids-AAA-34-36 | N/A |
| ORF8-AAA-37-39 | FL-ORF8 with triple Alanine substitution at amino acids-AAA-37-39 | N/A |
| ORF8-AAA-40-42 | FL-ORF8 with triple Alanine substitution at amino acids-AAA-40-42 | N/A |
| Rep-ORF8-WT | Spike deleted (replaced with eGFP-nLuc) Replicon (pBAC) plasmid of WA1 expressing ORF8 | N/A |
| Rep-ORF8-Stop | Spike deleted (replaced with eGFP-nLuc) Replicon (pBAC) plasmid of WA1 with ORF8 having stop codon at 2nd amino acid | N/A |
| IC-XBB.1.5 | Infectious clone (pBAC) plasmid of XBB.1.5 expressing truncated ORF8 (G8Stop) | N/A |
| IC-XBB.1.5+ORF8 | Infectious clone(pBAC) plasmid of XBB.1.5 expressing full length ORF8 | N/A |

### VLP production, purification, and confirmation

VLPs were produced by transfecting plasmids for all the structural proteins (S, N, M-IRES-E(MiE)) and ORF8 or EGFP, along with a plasmid expressing firefly luciferase with the SARS-CoV-2 RNA packaging signal(T-20), into HEK 293T cells, with a slight modification to a previously described protocol (21). The HEK293T cells were seeded in a 6-well with 0.5×10^6 cells/well. For transfection, S:N:MiE:T-20: ORF8/eGFP plasmids with 0.125:0.67:0.33:1:0.4 DNA mass ratios for a total of 2.525 µg of DNA were diluted in 150 µL OptiMEM. The X-treme Gene 9 (X9) DNA transfection reagent (Roche: 6365809001) was added 1:3 as DNA:X9 added to the plasmid dilution and mixed properly. This mixture was incubated for 15 minutes at room temperature and then added to cells. The culture media was replaced the next day. The culture media containing VLP collected 48 hours post-transfection, spun at 1000 rpm for 5 minutes, filtered through a 0.45 μm filter, and used immediately for infection or stored at -80°C for later infection. This filtered supernatant is considered VLP. For the experiment, where different ORF8 plasmid amounts were used, a vector plasmid was added to ensure the total ORF8+vector plasmid amount was 0.4μg. For infection, the HEK293T-ACE2-TMPRSS2 cells (50K cells/condition) were mixed with 100 μl of VLP and plated immediately in 96-Chimney opaque flat-bottom plates (Grenier). After 24 hours of incubation at 37°C, the cells were lysed with 30 μl/well of passive lysis buffer (E1941) for 15 minutes. After that, 50 μl of luciferase assay buffer (Promega, E1501) was added and luminescence was read using the TECAN plate reader (Infinite® 200 PRO) with auto attenuation, and 1000 ms integration.

To purify the VLPs, the transfection was done in 10 cm2 plates with the same plasmid ratio with total 24.125ug of plasmid and 72.4ul of PEI was used as transfection reagent. The filtered supernatant of producer cells (collected 48 hr after transfection) was combined with 0.5–1 ml of a control lentivirus, placed in an ultracentrifuge tube and a 20% sucrose solution (1/10 of total volume) was carefully added to the bottom of the tube. PBS was added to adjust volume and balance, and the tubes were spun in an SW32Ti at 24,000 rpm for 2 hours. The supernatant was decanted and the pellet dried for 10 mins. The pellet was then resuspended in 90 ul RIPA Buffer (25 mM Tris-HCl pH 7.6, 150 mM NaCl, 1% NP-40, 1% sodium deoxycholate, 0.1% SDS) and analyzed by western blot. The lentivirus served as a control for concentration. The control lentivirus was produced with a 1:1:0.34 DNA ratio with plasmids pLVX-EF1α-IRES-Puro: psPAX2: pMD2.6.

### Western blot analysis

Cells were collected by spinning at 1000 rpm for 5 minutes, followed by a 1x PBS wash. Cells were lysed in Flag lysis buffer (25 mM Tris-HCl pH 7.4, 150 mM NaCl, 1 mM EDTA, 1% NP-40, supplemented with Halt protease inhibitor cocktail). For infected cells or purified VLP, RIPA buffer was used to obtain whole-cell lysates. The protein concentration in the cell lysates was determined using a DC^TM^ protein assay kit (BioRad, 5000111). The same amounts of proteins were run on 4-20% Mini-PROTEAN® TGX™ Precast Protein Gels (BioRad, Cat:4561096) or homemade 6-15% SDS-PAGE gels and transferred to a nitrocellulose membrane (Biorad). The membrane was blocked in 10% non-fat dry milk in TBS-T and stained with primary antibody overnight at 4°C or 2-3 hours at room temperature. Blots were rinsed with TBS-T three times for 5 minutes each and stained with secondary HRP antibody (Bethyl A90-516P (mouse), A120-201P (rabbit) 1:5000). After that three washes were done with TBS-T. The blot was incubated with a chemiluminescence kit (Roche 12015200001, Thermo Scientific™ 34096) and images were captured using ChemiDoc™ Imaging System (Biorad 12003153). Densitometry was done using ImageJ software. All the antibodies used in this study are listed in Table 2.

**Table 2.**

| **Antibody** | **Manufacturer** | **Cat: No** |
| --- | --- | --- |
| SARS-CoV2 Spike(S2) | Abcam | ab272504 |
| SARS-CoV-2 Spike S1 Alexa 647 | R&Dsystems | FAB105403R |
| SARS-CoV2 Spike Alexa 594 | Novus Biologicals | NBP2-90980AF594 |
| SARS-CoV2 Nucleocapsid (N) | Abcam | ab273434 |
| SARS-CoV2 Nucleocapsid (N) | Sino-biological | 40143-MM05 |
| SARS-CoV2 ORF8 | Abcam | ab283914 |
| Anti-HIV-1 P24 | Sigma Aldrich | SAB3500946 |
| Strep Tag II | Qiagen | 1023944 |
| Flag | Sigma Aldrich | F3165 |
| hACE2 | Proteintech | 28868-1-AP |
| hTMPRSS2 | Abcam | ab109131 |
| GAPDH | Cell Signaling | 5174S |
| GAPDH | Santa Cruz | sc-365062 |
| GM130- Alexa Fluor™ 555 | ThermoFisher Scientific | PA1-077-A555 |
| Hoescht 33342 | Invitrogen | H3570 |
| Mouse IgG-AlexaFluor549 | Invitrogen | A11005 |
| Rabbit IgG-AlexaFluor488 | Invitrogen | A11008 |
| Rabbit IgG-HRP | Bethyl | A120-201P |
| Mouse IgG-HRP | Bethyl | A90-516P |

### Immunoprecipitation

Transfected HEK293T cells were collected at the 48 hr timepoint and lysed in IP buffer (50 mM Tris-HCl, 150 mM NaCl, 1 mM EDTA, 1% NP40, supplemented with Halt protease inhibitor cocktail). For pulldown, 0.5-1 mg of lysate was incubated overnight with 30 ul of Strep-Tactin Sepharose resin (IBA Life Science, 2-1201-002), rotating at 4°C. Bound protein was washed five times with IP buffer and eluted with Strep-Tactin elution buffer (IBA Life Sciences, 2-1000-025). Eluted samples were analyzed by western blot.

### Immunofluorescence microscopy

A549-ACE2 cells were infected with MOI 0.5 and plated onto 22 mm^2^ no. 1.5 coverslips. Cells were fixed in 4% paraformaldehyde, permeabilized with methanol on ice for 10 min, and blocked in 3% bovine serum albumin. Cells were then immunostained with the antibodies indicated in Table 2. The coverslips were mounted onto glass slides using ProLong Gold Antifade Mountant (Invitrogen, P36934) and analyzed by confocal microscopy (Olympus FV3000RS) using an Olympus UPLAN S-APO 60× OIL OBJ,NA1.35,WD0.15MM objective. The resulting Z-stack was reconstructed and rendered in 3D using Imaris software (Oxford Instruments).

### VLP entry assay with recombinant ORF8

Flag-tagged ORF8 (conc. 1.7mg/ml) was produced by Chempartner (Batch: CP20210318-MA, diluted in a buffer containing 50 mM Tris, 200mM NaCl, 1mM DTT, pH 8.0). We tested 2 concentrations 1ug/ml and 10ug/ml. Three independent batches of VLPs were produced without ORF8, and 100 ul of each batch was incubated with recombinant ORF8 for 1.5 hr at 37°C. This was used to infect HEK293T-ACE2-TMPRSS2 cells that had been plated the day before. 24 hr after infection, cell viability and infection was measured using Cell titre Glo assay (Promega, G7571), and Luciferase assay system (Promega, E1501).

### Quantification and statistical analysis

The number of experiments and replicates are indicated in individual figure legends. Data was processed and visualized using GraphPad Prism. All quantified data are represented as mean ± SD, as indicated, and quantification details are available in figure legends. Western blot band intensities were quantified using ImageJ.

## Data availability

All data supporting the present study’s findings are available in the article, extended data and supplementary figures, or from the corresponding authors on request. Source data are provided with this paper.

## Acknowledgments

M.O. received support from the National Institutes of Health (U19AI135990), the James B. Pendleton Charitable Trust, the Roddenberry Foundation, P. and E. Taft, and the Gladstone Institutes. M.O. is a Chan Zuckerberg Biohub – San Francisco Investigator. M.O. also thanks Fast Grants and the Innovative Genomics Institute for their support. J.A.D. acknowledges support from the National Institutes of Health (R21AI59666) and from the Howard Hughes Medical Institute and the Gladstone Institutes. I.P.C. was supported by NIH/NIAID (F31 AI164671-01). M.M.K. and M.O. highly acknowledge senior scientific editor Françoise Chanut for her valuable input in editing the manuscript. We thank Chia-Lin Tsou and Tan Yee Joo for helping with shipment logistics for the Singapore isolate (EPI_ISL_414378).

## Author contributions

Conceptualization: M.M.K., M.O.; Methodology and Investigation, M.M.K., I.P.C, F.W.S, T.Y.T., T.T., R.K.S., A,M.S., A.C., M.M.M., U.S.G., J.H., I.J.K., J.B.; Writing, M.M.K., M.O.; Analysis, M.M.K., I.P.C., Visualization, M.M.K., Isolate acquisition: S.W.F., G.R.K, L.R., L.F.P.N., Supervision Funding Acquisition, N.J.K., J.A.D., E.V., M.O.

