## Supplemental Figures for "Regulation of virion production by the ORF8 signal peptide across SARS-CoV-2 variants"

### Supplementary Figures

**Supplementary figure S1:** Conservation of ORF8 mutations in the SARS-CoV-2 B lineage. The phylogeny shows relationships in different SARS-CoV-2 clades (not based on ORF8 mutations) as defined by Nextstrain (38). The phylogeny was adapted from CoVariants (39). Each new ORF8 mutation is placed in the variant where it first appeared, and the colored line shows which mutation continued to subsequence lineages. The mutations that did not pass on to major lineages or sub-lineages are colored as grey. The mutations that passed on to next colored either dark cyan (S84L) or Cadmium Orange (G8*). * = stop codon in this image.

**Supplementary figure S2 (related to figure 1): A.** Protein expression in HEK-293T-A/T cells infected by clinical isolate. Spike shows two bands, top band is the uncleaved version(S0, ~180 kDa), the next prominent band close to ~90kDa is S2. **B.** Representative western blot of BHK-21 cells transfected with replicon plasmids and a plasmid encoding SARS-CoV-2 N (Delta variant). n = 3 independent experiments.

**Supplementary figure S3 (related to figure 2):** A549-ACE2 cells were infected with WA1 and Singapore clinical isolates. Immunostaining was done with ORF8 (Green), Spike (Turquoise), and GM130 (Red; cis-Golgi marker) antibodies. DAPI (Blue) staining was used for nucleus. White dotted square area was enhanced three times. White bar = 8 μm. n = 3 independent experiments.

**Supplementary figure S4 (related to figure 3) :** SARS-CoV-2 ORF8 does not affect VLP entry but affects VLP production. **A.** Normalized luminescence of VLP-infected cells. VLP was incubated with different amount of recombinant ORF8 prior to infection.

Cell viability was not affected by VLP infection. Three different stocks of VLPs were used in this experiment. **B-D.** Normalized luminescence of VLP-infected cells. VLPs were produced in the presence of a constant amount of eGFP or ORF8 plasmid and an increasing concentration (1-fold, 2-fold and 4-fold) of each structural protein plasmid. Data are represented as means plus SD. For each condition, three technical replicates were done for luminescence measurements with total 3 independent experiments were done. The P values were calculated by One-way ANOVA test. *, $P \leq 0.05$; **, $P \leq 0.01$; ***, $P \leq 0.001$; ns, not significant. n = 3 independent experiments.

**Supplementary figure S5:** SARS-CoV-2 ORF8 affects Delta and Omicron VLP production. **A-B**. Normalized luminescence of cells infected with B.1, Delta, and Omicron VLPs. **A.** VLPs were produced from the respective variants' structural proteins, in the presence of ORF8 or control plasmid (eGFP). **B.** VLPs were made using Spike from either the Delta or Omicron variant, and other structural proteins (E, M and N) from B.1. For each condition, three technical replicates were done for luminescence measurements with total 3 independent experiments were done. **C-D.** Representative western blot of VLP-producing cells (from **A** and **B**). Data are represented as means plus SD. The P values were calculated by One-way ANOVA test. *, $P \leq 0.05$; **, $P \leq 0.01$; ***, $P \leq 0.001$; ns, not significant. n = 3 independent experiments.

**Supplementary figure S6 :** SARS-CoV-2 ORF8 disulfide bonds have no role in VLP production. **A.** Schematic diagram of cysteine residues in ORF8. C20 mediates ORF8 dimerization. The other 6 cysteine residues forms 3 intra-molecular disulfide bonds. **B,D.** Normalized luminescence of VLP-infected cells. VLPs were produced in the presence of different ORF8 cysteine mutants. For each condition, three technical replicates were done for luminescence measurements with total 3 independent experiments were done.ORF8-NoCys is a ORF8 construct with all cysteine residue converted to alanine **C,E**. Representative western blot of VLP-producing cells (from B, D). Data are represented as means plus SD. The P values were calculated by One-way ANOVA test. *, P ≤ 0.05; **, P ≤ 0.01; ***, P ≤ 0.001; ns, not significant. n = 3 independent experiments.

**Supplementary Figure S7 (related to Figure 4):** The first 40 aa of ORF8 are necessary and sufficient to bind Spike and significantly reduce VLP production. **A, B.** Representative western blots of protein expression and immunoprecipitation of the cell lysate of VLP-producing cells (related to **Figure 4A-B**). VLPs were produced in the presence of different fragments of ORF8. n = 3 independent experiments.

**Supplementary figure S8 (related to figure 5):** Impact of triple alanine mutant scan of ORF8 from aa 16 to aa 42 on VLP production. **A.** Schematic diagram of triple alanine scan. FL = full length. A = alanine. **B.** Normalized luminescence of VLP-infected cells. VLPs were produced in the presence of either ORF8 (ORF8-FL or AAA mutants) or eGFP. The supernatant of VLP-producing cells was used to infect HEK293T-A/T. Luminescence was normalized to eGFP controls for each experiment. For each condition, three technical replicates were done for luminescence measurements with total 3 independent experiments were done. **C**. Representative western blot of VLP-producing cells (from **B**) showing protein expression and immunoprecipitation. Data are represented as means plus SD. The P values were calculated by One-way ANOVA test. *, P ≤ 0.05; **, P ≤ 0.01; ***, P ≤ 0.001; ns, not significant. n = 3 independent experiments.

**Supplementary Figure S9 (related to Figure 5):** A549-ACE2 cells were infected with XBB.1.5 recombinant viruses. Immunostaining was done with ORF8 (Green), Spike (Turquoise) and GM130 (Red; cis-Golgi marker) antibodies. DAPI staining was used for the nucleus. The white dotted square area was enhanced three times. White bar = 8 µm. n = 3 independent experiments.

Supplementary figure S1

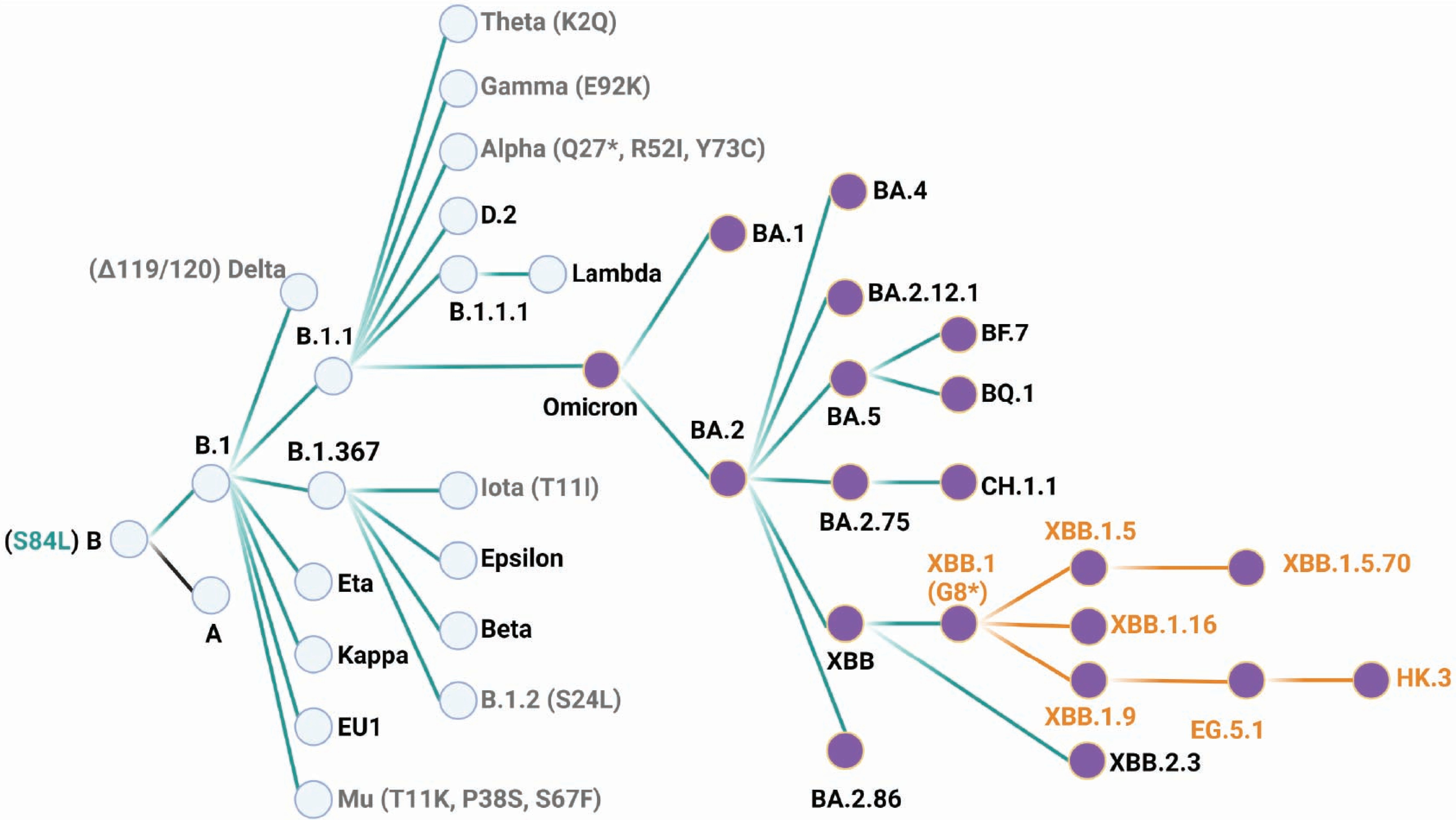

Supplementary figure S2

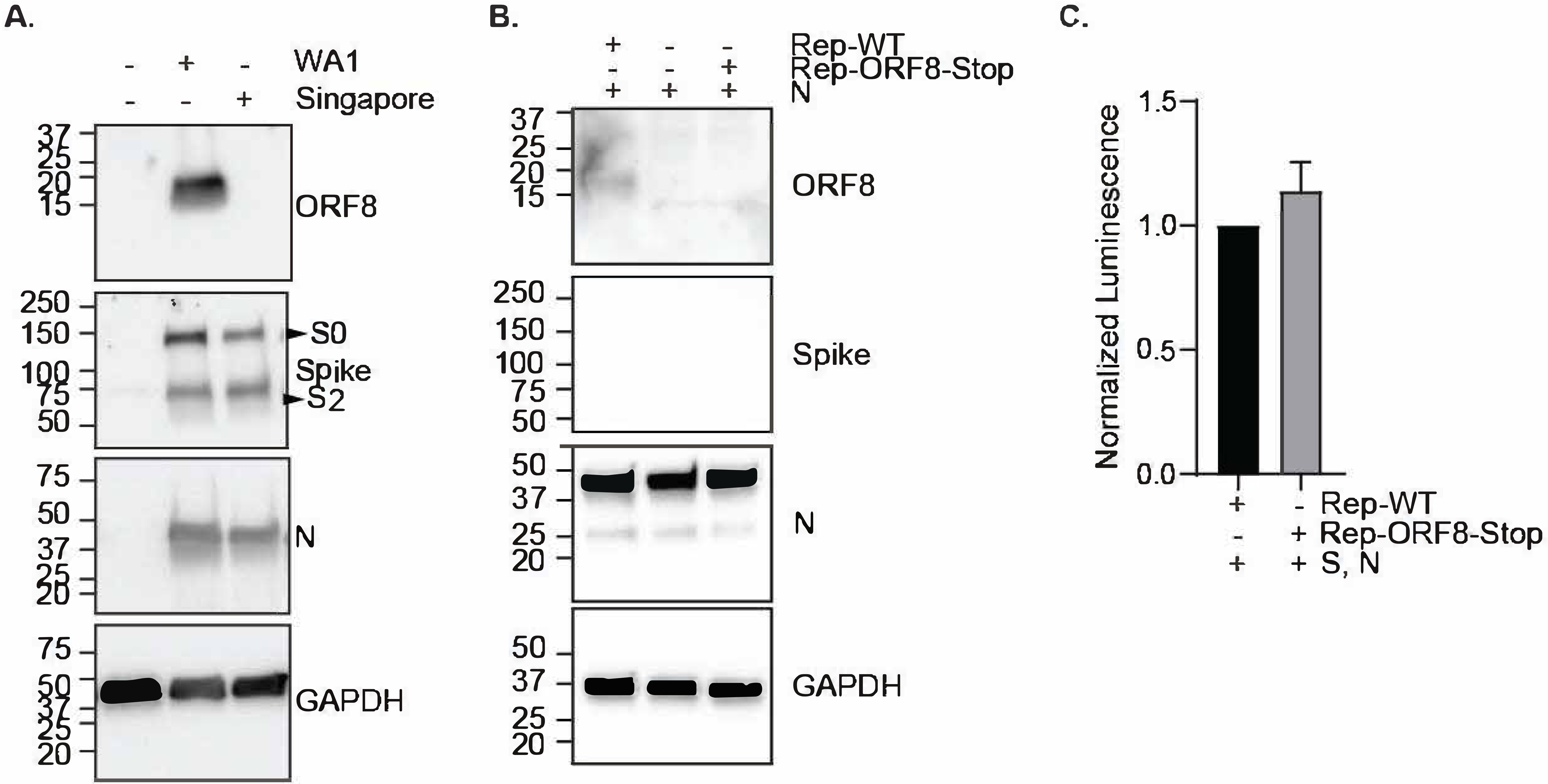

Supplementary figure S3

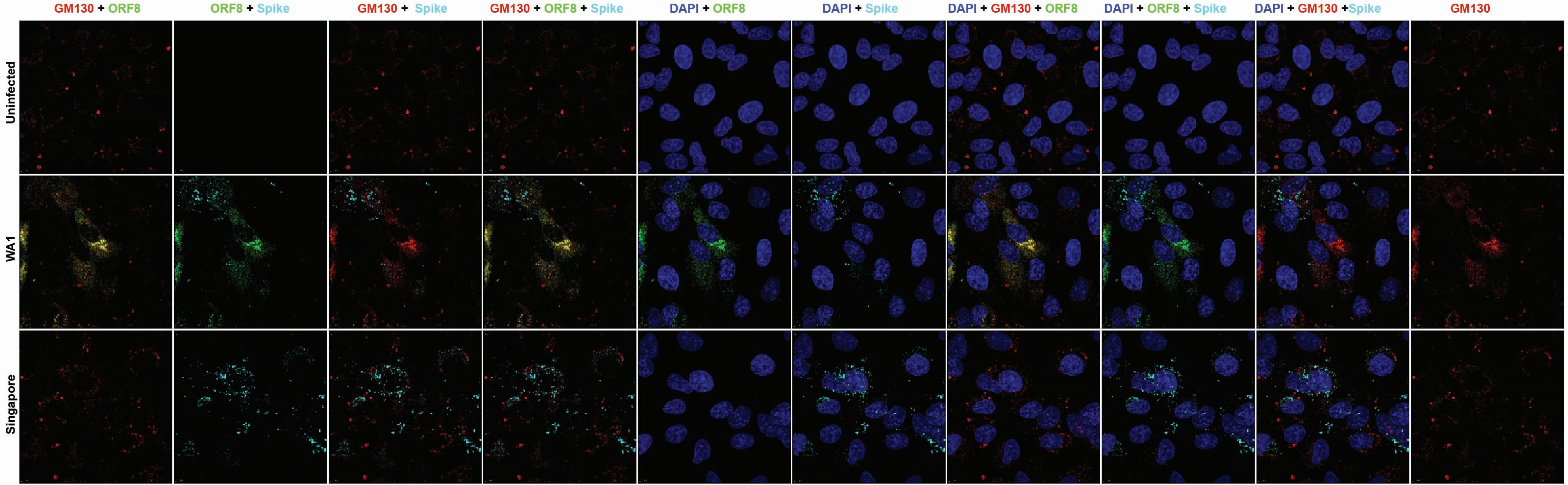

Supplementary figure S4

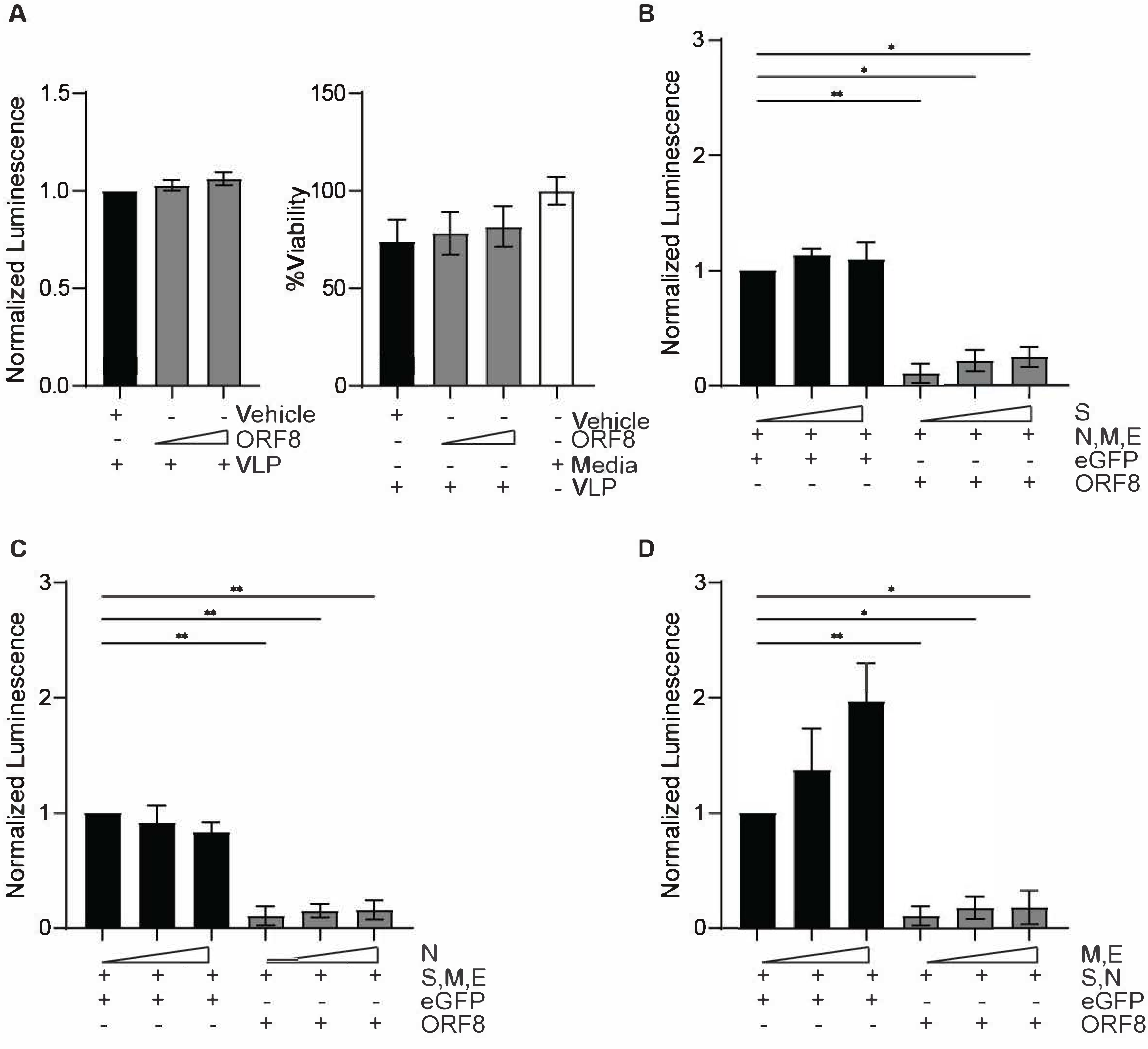

Supplementary figure S5

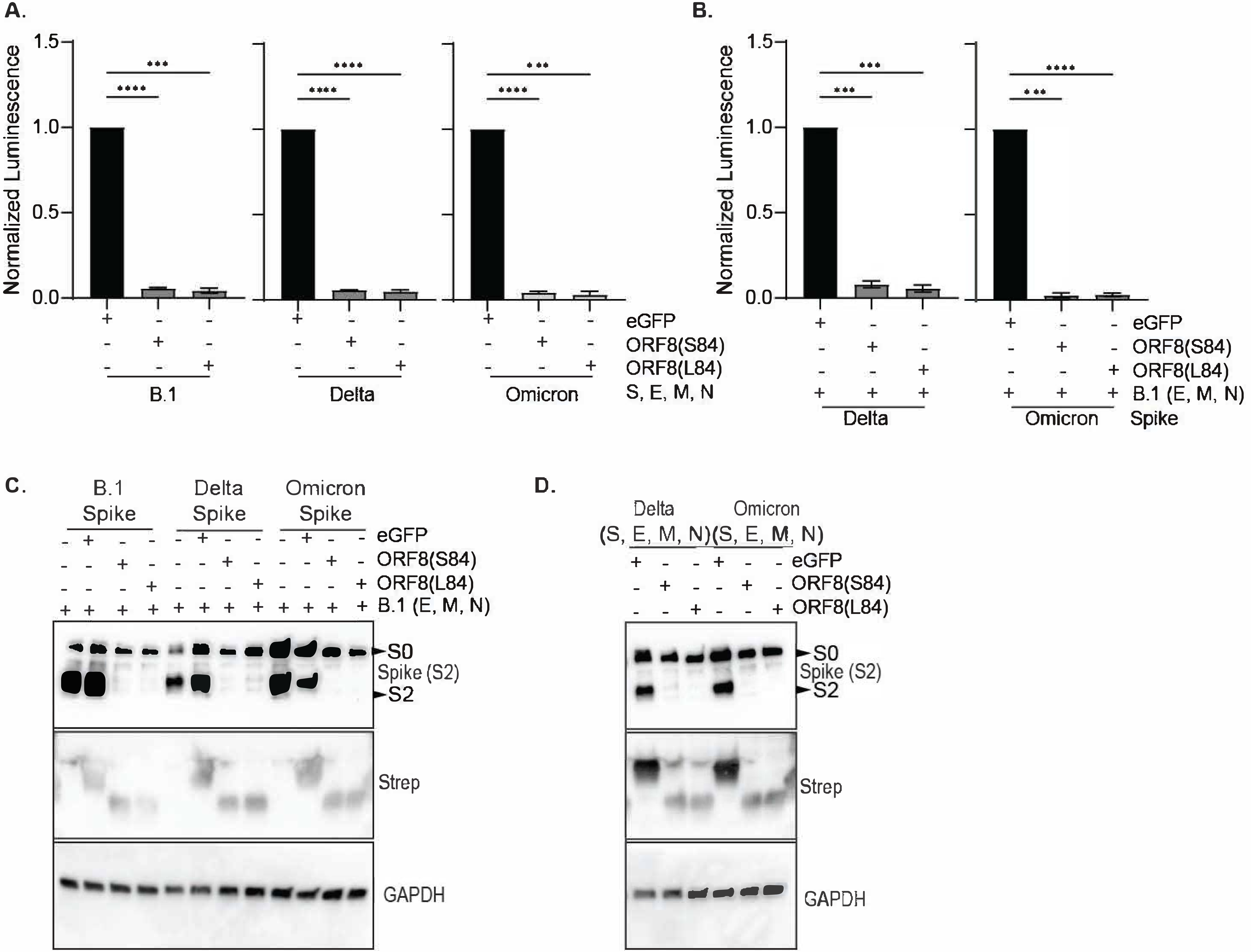

Supplementary figure S6

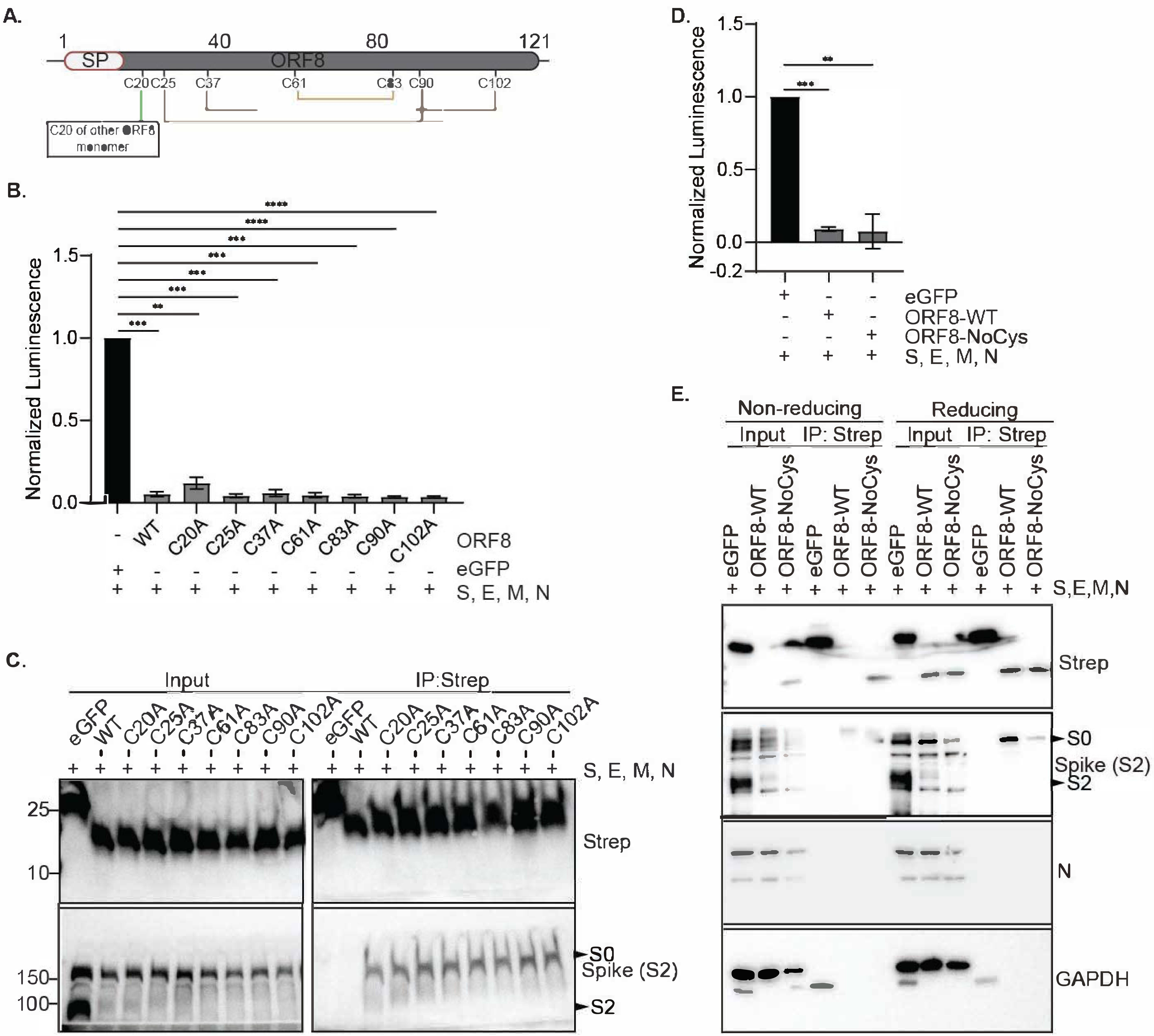

**Supplementary figure S7**

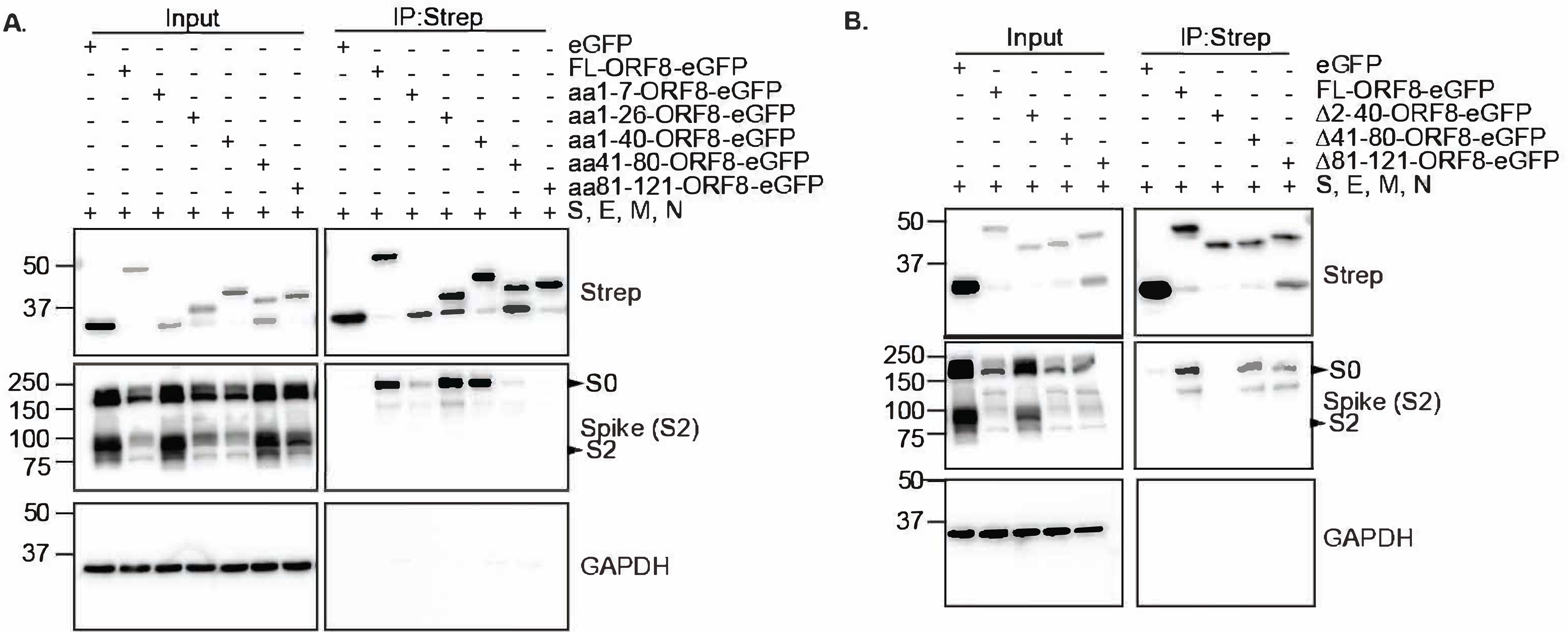

**Supplementary figure S8**

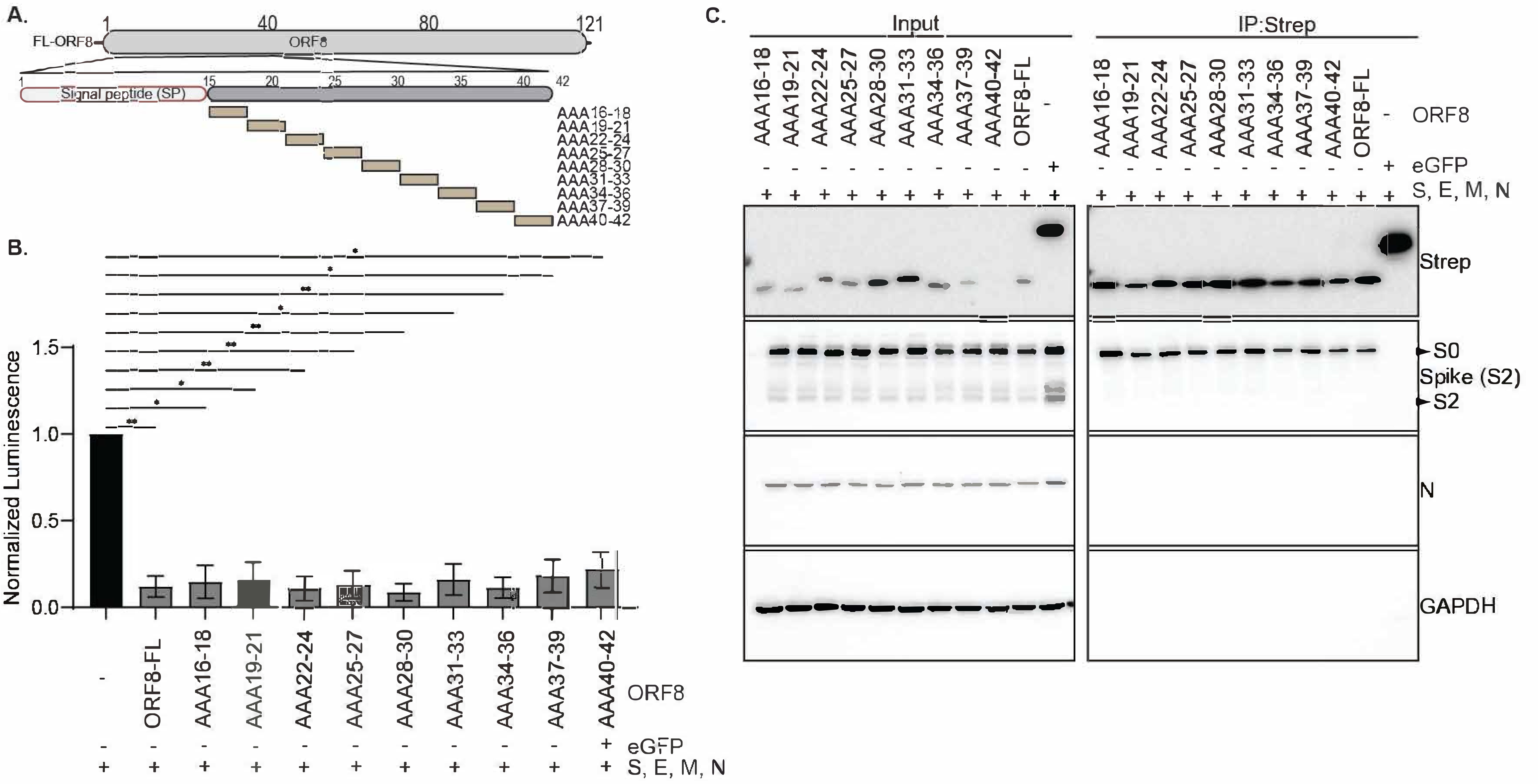

Supplementary figure S9

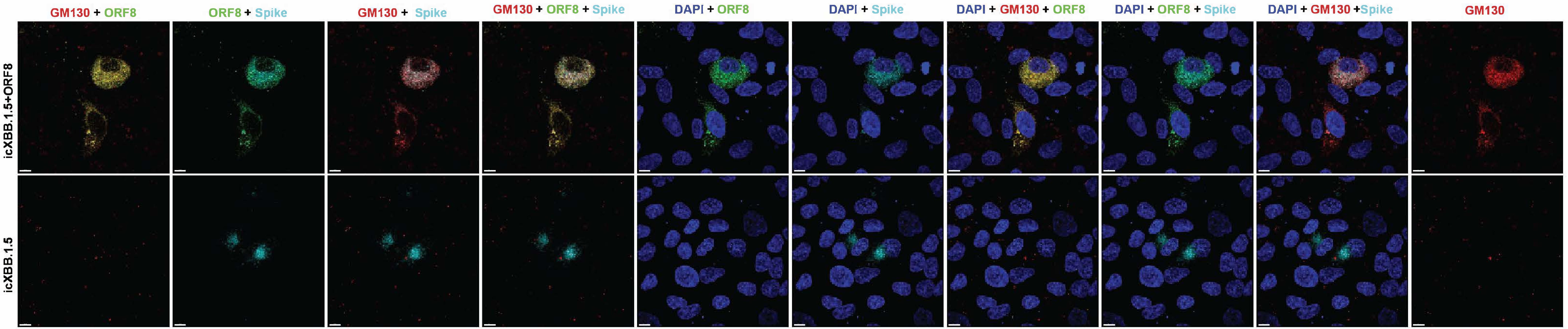
